# Effect of Aeromonas hydrophila infected on metabolomic response and gut microbial 16S rRna of *Charybdis japonica*

**DOI:** 10.1101/2021.10.25.465830

**Authors:** Mingming Han, Chenxi Zhu, Zakaria Zuraini, Tianheng Gao, Ying Yang, Tongqing Zhang, Feng Ji, Qichen Jiang

**Affiliations:** Freshwater Fisheries Research Institute of Jiangsu Province. 79 Chating East Street, Nanjing 210017, China; Biology Program, School of Distance Education, Universiti Sains Malaysia, 11800 Minden, Penang, Malaysia; Institute of Marine Biology, College of Oceanography, Hohai University, China; Jiangsu Key Laboratory for Molecular and Medical Biotechnology, College of Life Sciences, Nanjing Normal University, Nanjing, 210023, China

**Keywords:** Metabolome, innate immunity, 16S rRNA, *Charybdis japonica*, *Aeromonas hydrophila*

## Abstract

The innate immune response of *Charybdis japonica* treated with *Aeromonas hydrophila* was explored using bioinformatics. Metabolomics data were integrated with a gut microbial 16S rRNA dataset, together with information on corresponding enzyme activity. The results of the study showed that after being infected with *A. hydrophila*, some beneficial genera of bacteria in the gut of *C. japonica*, such as Photobacterium, Rhodobacter, Polaribacter, Psychrilyobacter, Mesoflavibacter, Fusibacter and Phormidium, could directly inhibit Vibrio or produce extracellular polysaccharides with highly effective antibacterial properties. The intestinal probiotics of *C. japonica* such as Mesoflavibacter have a mutually reinforcing relationship with Phaeobacter, Colwellia, Bacillus, Psychrobacter and Cohaesibacter. Conditional pathogenic bacteria in the gut of healthy crabs may also have such a symbiotic relationship with intestinal probiotics, promoting their growth and reproduction. For example, Phormidium has a mutualistic relationship with *Aeromonas* and Azopira. Metabolites in the gut of *C. japonica* infected with *A. hydrophila*, including beta-alanine metabolism, nitrogen metabolism, inositol phosphate metabolism, galactose metabolism, histidine metabolism, ascorbate and arginine and proline metabolism were increased, with alanine metabolism being the most abundant. The activity of metabolite related enzymes such as lipid peroxidase, phenoloxidase, superoxide dismutase, nitric oxide synthase, glutathione transferase and mid-glutathione decreased and NO levels also decreased. The positive correlation with the probiotic flora suggests that metabolites increase with bacterial abundance and that microbial metabolites or co-metabolites can, in turn, achieve many pleiotropic effects to resist invasion by *A. hydrophila*. These results may contribute to further research in the resistance of *C. japonica* to invading pathogens.

**Importance:** With the rapid development of the *C. japonica* farming industry, investors, in pursuit of economic benefits, have encountered problems such as frequent outbreaks of various diseases, resulting in high mortality and huge economic losses. The open water circulation system can give rise to several crab bacterial diseases. Among these, *A. hydrophila* is a pathogenic bacterium affecting fish and crustaceans, which leads to huge economic losses. Our results suggest that metabolites increased with the abundance of bacteria. It is possible that the autoimmune system and the entry of *A. hydrophila* into the intestinal tissues of *C. japonica* react immunologically and that the organism is producing certain metabolites which may be metabolised by various bacteria and absorbed into the circulation. In addition, some of these metabolites are modified or bound in the hepatopancreas to produce microbiota-host co-metabolites. These microbial metabolites or co-metabolites can resist invasion by *A. hydrophila*.

## 1. Introduction

*Charybdis japonica* is found as an important edible species in the intertidal zone along the west coast of the Pacific Ocean, including China, Japan and Malaysia. The habitat of *C. japonica* is frequently affected by pathogenic bacteria of the Vibrio family(1). *A. hydrophila* belongs to the family Vibrionaceae and the genus *Aeromonas*, which is a group of thermophilic hydrophilic *Aeromonas*. It is ubiquitous in nature, is pathogenic to aquatic animals (2) and can cause septicaemia in various aquatic animals, bringing economic losses to the freshwater aquaculture industry, thus attracting the attention of fisheries, veterinary and medical researchers (3). In aquaculture, a complex micro-ecosystem exists in the gut that functions for digestion, nutrient absorption and disease resistance (4). The gut-associated microbiota plays a unique role in host gut development, immune response, disease resistance and homeostasis. Beneficial strains of bacteria and beneficial metabolites have been introduced into the host gut with significant results(5). The microbial composition of the gut of crustaceans has a significant impact on animal health, growth and survival. Factors such as the internal structure of shrimps and crabs, host conditions, diet, climate change, living environment, and bacterial or viral infections can all affect the gut flora of aquatic animals(5). This experiment is designed to understand the role of bacteria in the gut of crabs, firstly by understanding the composition of host gut bacterial flora. Secondly, it is important to discover the changes in the composition of the gut bacterial flora after infection with the bacteria.

For crabs, the microbial diversity of crab carapace, gut and haemolymph fluid was studied by using 16S rRNA gene analysis, cloning and sequencing (6), and various potential pathogens were examined, such as *Alternaria alternata*, *Bacillus*, *Escherichia coli*, photobacterium subspecies and *Vibrio harveyi*. They have greatly expanded our view of the microbial life associated with marine invertebrates, such as microbial community composition, functional potential and metabolic activity (7). In recent years, the diversity of gut microbes in a variety of aquatic animals has been studied based on 16S rRNA genes, and the gut microbial diversity of shrimp has been investigated by molecular isolation (8). A comparison of the gut microbiota of healthy and diseased crabs revealed that the relative abundance of bacteria in healthy crabs was higher than in diseased crabs (9). *Scylla paramamosain* was infected with *Vibrio vulnificus* and strains were screened for antagonistic activity against *Vibrio parahaemolyticus* using an agar spot assay. The antagonistic strains were then identified by 16S rRNA gene sequence analysis (10). The 16SrRNA method has been important in the analysis of crab infection with bacteria.

Metabolomics is a method of histological measurement and an important tool in histology. Metabolomics methods allow the unbiased analysis of the composition of all detectable metabolites, and thus, rapid quantitative detection of stress responses(11). Metabolomics has been shown to provide in-depth research data in crab diet, climate change, living environment, physiology and pathology and biochemistry (12). *A. hydrophila* infects aquatic animals through incidental bruises on the body. Metabolomics is also an important tool to study the innate immunity of *C. japonica*.

This experiment first studied the innate immunity of *C. japonica* infected with *A. hydrophila*. The aims of the study were 1) to examine the composition of the intestinal bacterial flora by the 16S rRNA sequencing technique, discover the changes in the composition of the intestinal bacterial flora after infection with *A. hydrophila*; 2) to find differential metabolites and the function of these metabolites through metabolome sequencing; 3) to investigate and understand the relationship between metabolites and gut microbes by integrating metabolomics data with gut microbial 16S rRNA datasets using bioinformatics tools; 4) to analyse relevant enzyme activities that affect metabolites.

## 2. Experimental methods

### 2.1. Experimental materials

Samples of *C. japonica* were collected from the South China Sea. The crabs weighed 81 ± 3.4 g and body length and width were 8 ± 1.7 cm and 6 ± 1.6 cm, respectively. One hundred and twenty *C. japonica* were divided equally into six aquariums with identical nets and PVC pipes to act as a shelter for the animals and prevent cannibalism. They were fed (9812; Shanghai Harmony Feed Co., Ltd., China) at 8:00 and 18:00 daily with the equivalent of 5% of the body weight of a pair of *C. japonica*. The crabs were exposed to seawater using an artificial sea salt cycle (salinity 28 psu), with water temperature controlled at 25 ± 1°C, pH 8.0 ± 0.2, a dissolved oxygen concentration of 5.0 mg L^−1^ and a 12 h light/dark cycle.

After 2 weeks of acclimation, water quality was maintained at the same level as in the acclimation period. Animals were not fed commercial feed for 24 hours before the trial. Sixty animals were removed from three aquaria for bacterial infection experiments, and 10^5^ CFU/L of *A. hydrophila* was injected into the fourth leg. An equal amount of saline was injected into the fourth leg of 60 animals in the remaining three aquaria, and samples were collected 24 hours later. Intestinal and hepatopancreas tissue was collected with sterile scissors and forceps after anaesthetizing the crabs on ice. All intestinal tissues and hepatopancreas were then rapidly frozen in liquid nitrogen and stored at −80°C for subsequent experiments. To avoid errors due to individual differences, composite samples were prepared by mixing equal amounts of each group of intestinal tissues and then aliquoting them into two samples. Some samples were assayed for relevant immune genes and enzymes. The *A. hydrophila* for this study was provided by Shanghai Ocean University. *A. hydrophila* freezing tubes were removed from storage at −80°C and quickly transferred to a water bath at 37°C to rapidly dissolve the bacterial freezing tract solution. After observation, to ensure that *A. hydrophila* was free of contamination, LB solid plates were recoated with bacteria and incubated in a constant temperature incubator at 28°C for 20 h. The *A. hydrophila* were grown to log phase and diluted to 10^5^ cfu/L by the McElloby method.

### 2.2. DNA extraction and PCR amplification

The mixed intestinal tissues were subjected to total DNA extraction using an E.Z.N.A.® soil kit (Omega Bio-Tek, Norcross, GA, USA). DNA concentration and purity were assayed using NanoDrop2000. DNA extraction quality was checked by 1% agarose gel electrophoresis. PCR amplification of V3-V4 was performed using primers 338F (5 ′ -ACTCCTACGGGAGGCAGCAG-3 ′) and 806R (5 ′ -GGACTACHVGGGTWTCTAAT-3 ′). PCR products were recovered using a 2% agarose gel, and purified using the AxyPrep DNA Gel Extraction Kit (Axygen Biosciences, Union City, CA, USA), eluted in Tris-HCl and detected by 2% agarose electrophoresis. Quantification was carried out by using QuantiFluor™-ST (Promega, USA).

### 2.3. Illumina Miseq sequencing

The purified amplicons were constructed into PE 2*300 libraries according to the Illumina MiSeq platform (Illumina, San Diego, USA) standard operating protocols. The raw sequence fastq files were imported into a file format ready for subsequent processing by QIIME2. The QIIME2 dada2 plug-in was then applied for quality control, pruning, denoising, splicing and removal of chimaeras to obtain the final table of characteristic sequences. Next, the ASV representative sequences were compared to the 99% similarity GREENGENES database (the database was trimmed to the V3V4 region based on the 338F/806R primer pair) to obtain a taxonomic information table for the species, removing all contaminating mitochondria. Methods such as ANCOM, ANOVA, Kruskal Wallis, igraph, LEfSe and DEseq2 were used to identify groupings. The R package “mixOmics” was used for bacteria with differences in abundance between samples, and partial least squares discriminant analysis (PLS-DA) was used as a supervised discriminant analysis statistical method to reveal the relationship between microbial communities and sample classes and to enable the prediction of sample classes based on the relative abundance of the main microbial species in the sample. We also used co-occurrence analysis to calculate Spearman’s rank correlation coefficients, which were used to understand associations between species. In addition, PICRUSt software was used to predict the likely functional composition of the microbial community. Unless specifically noted, the parameters used in the above analyses are the default settings.

### 2.4. Metabolite extraction

Intestinal tissue (100 mg) was placed into a 5 mL tube (reduced by equal proportions if the sample size was insufficient) and mixed thoroughly for 1 min with 500 μL ddH_2_O at 4 °C. Methanol (1 mL, −20 °C) and heptadecanoic acid (60 μL 0.2 mg/mL) were added, shaken in a Vortex for 30 s, sonicated for 10 min at room temperature, left on ice for 30 min and centrifuged at 12 000 *g* for 10 min at 4 °C. The supernatant was transferred to a new 1.5 mL centrifuge tube and the sample was concentrated by vacuum centrifugal concentrator. Methoxy solution (60 μL) was added and shaken in a Vortex for 30 s. The reaction was carried out at 37 °C for 2 h. Finally, 60 μL of reagent (containing 1% trimethylchlorosilane) was added and the reaction was carried out at 37 °C for 90 min and centrifuged at 10 000 g for 10 min at 4 °C. The reaction was carried out at 37 °C for 90 min and centrifuged at 10 000 g for 10 min at 4 °C. The supernatant was added to a vial and part was used in quality control to correct for deviations in the results of mixed samples and errors caused by the analytical instrument, the remaining sample was used for GC-MS.

### 2.5. On-board testing

Gas chromatography was performed on an HP-5MS capillary column (5% benzene/95% methyl polysiloxane 30 m × 250 μm i.d., 0.25 μm film thickness, Agilent J & W Scientific, Folsom, CA, USA) with a constant flow of helium at 1 mL/min. The 1 μL sample was injected through an autosampler at a 20:1 split ratio. The injection temperature was 280 °C, set to 150 °C for the interface and adjusted to 230 °C for the ion source. Mass spectrometry was performed using a full scan method with a range of 35 to 750 (m/z)

### 2.6. Data pre-processing steps

The data obtained were subjected to GC-MS metabolomics assays for bioinformatics analysis. The raw data were converted into netCDF format (xcms input file format) by an Agilent MSD ChemStation workstation for peak identification, peak filtration and peak alignment. A data matrix including mass to charge ratio (m/z) and retention time and peak area (intensity) was obtained. The metabolites were annotated in conjunction with the AMDIS program using the National Institute of Standards and Technology (NIST) commercial database and the Wiley Registry Metabolome database. The metabolite alkane retention indices were provided according to The Golm Metabolome Database (GMD) (http://gmd.mpimp-golm.mpg.de/) and used for further substance characterisation. The detailed data results, which are normalised internally to allow for comparison between different quantities of data.

### 2.7. Measuring the activity of hepatopancreas-related enzymes

Hepatopancreas (100 mg) was homogenised with nine times the volume of physiological saline solution by weight to make a 10% tissue homogenate. The homogenate was centrifuged at 2500 rpm at 4°C for 10 min and the supernatant was placed on ice for assays. The activities of hepatopancreas-related enzymes were measured using kits from the Nanjing Jiancheng Company. Protein concentrations were determined by the Folin-phenol method and hexokinase activity was determined by a colourimetric method coupled to 6-phosphoglucose dehydrogenase. Superoxide dismutase (SOD), phosphofructokinase (PFK), pyruvate kinase (PK), nitric oxide synthase (i-NOS), lipid peroxidase (LPO), phenoloxidase (POX), glutathione transferase (GST), glutathione (GSH), glutathione peroxidase (GPX) and glutathione reductase (GR) were tested according to the manufacturer’s instructions. GPX was measured at 412 nm absorbance, GSH at 405 nm absorbance and GST at 412 nm absorbance. GST was measured by absorbance at 340 nm using 1-chloro-2,4-dinitrobenzene as the substrate. The absorbance of each tube was measured at 550 nm with a 1 cm optical diameter of 0.4 mm inner diameter and blanked with distilled water to detect nitric oxide (NO).

## 3. Results

### 3.1. *C. japonica* 16S rRNA sequencing data

After removing low-quality reads, a total of 577,767 valid reads were obtained from the 12 samples. The highest number of valid reads in all the samples of AH1 was 58,952 (Figure 2A) and the lowest number of valid reads in all the samples of CK was 32,453. The mean value of valid reads was 48,147.25. The highest number of optimized non-chimeric sequences in all the samples of AH1 was 42540 and the lowest number of optimized non-chimeric sequences in CK6 was 14226. The abundance of microorganisms in the gut of *C. japonica* can be assigned to the most classifiable taxa such as phylum, phylum, order, family and genus. Of the total 48 samples from the three groups, 99% of the phylotypes belonged to only four core phyla: Proteobacteria (0.92%), Tenericutes (5.21%), Fusobacteria (2.61%) and Bacteroidetes (0.73%; Figure 2).

**Figure 1:**
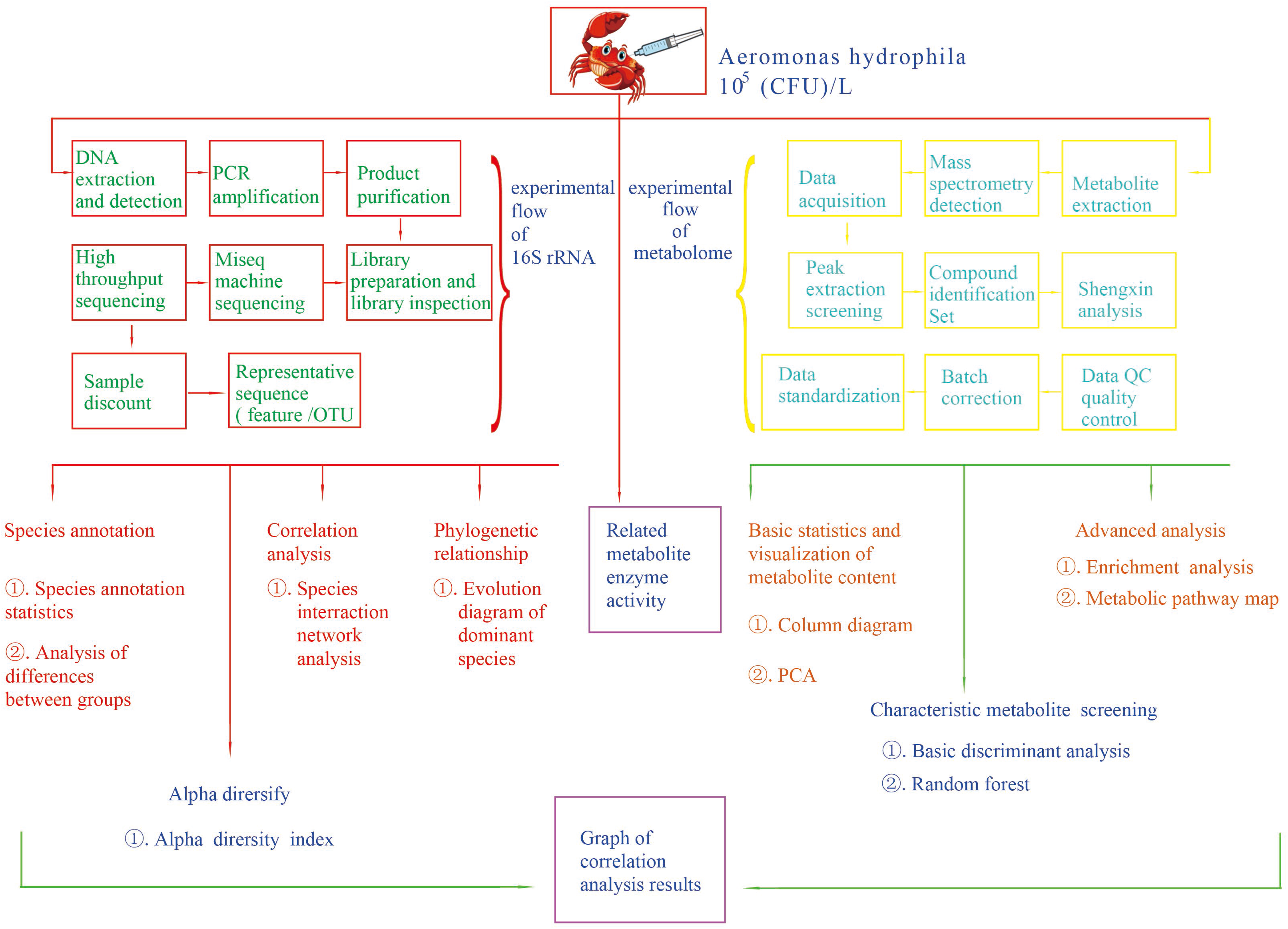
Workflow for the analysis of 16SRNA and metabolomic data in response to berberine.

**Figure 2:**
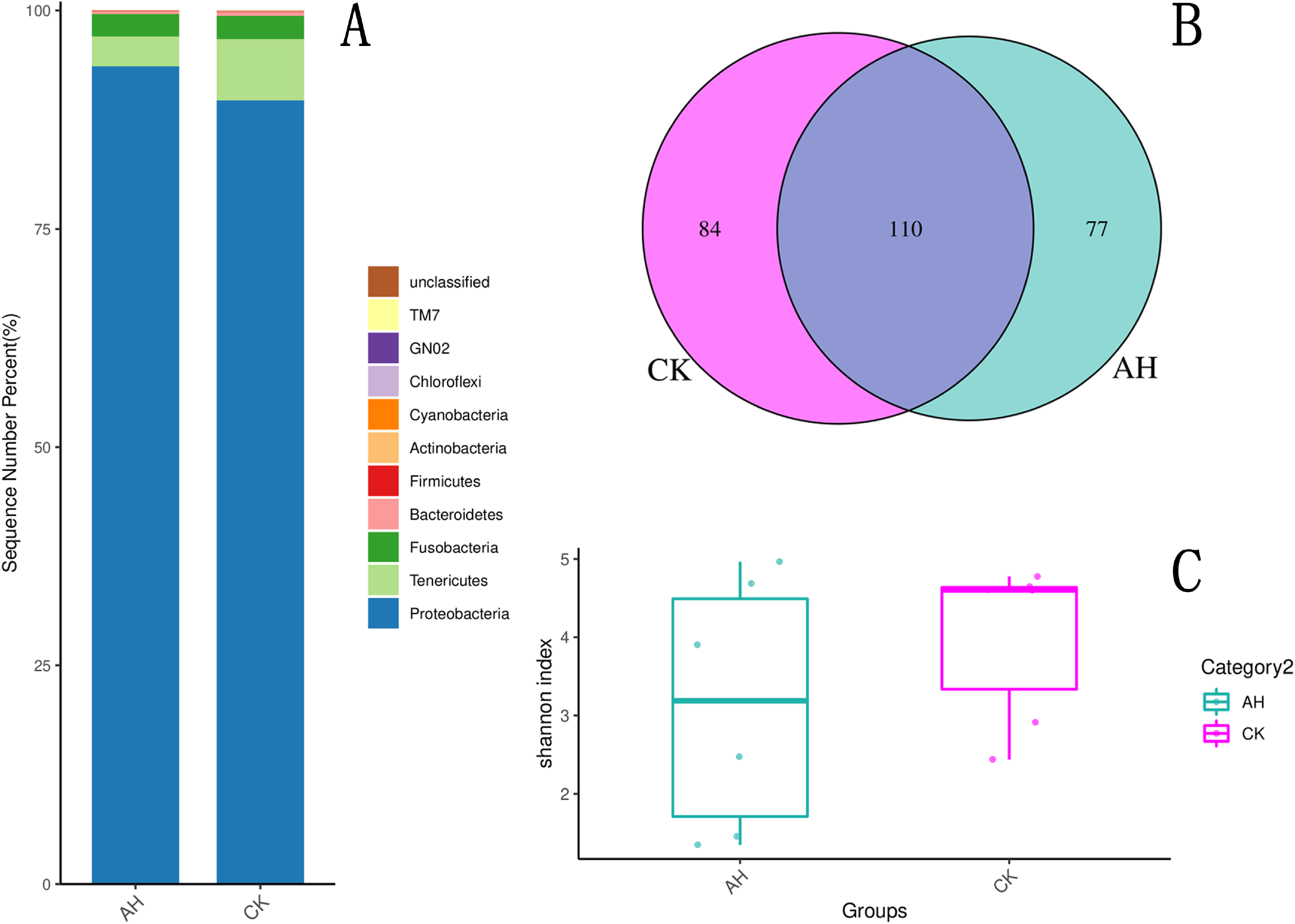
(A) Histogram of the relative distribution of each subgroup at the phylum level (top 15 species in relative abundance), the vertical coordinate (sequence number percent) indicates the ratio of the number of sequences annotated to that phylum level to the total annotated data, the top-down colour order of the histogram corresponds to the colour order of the legend on the right. *C. japonica* injected with 10^5^ CFU/L of *A. hydrophila* (AH) and control (CK). (B) Venn diagram of shared or endemic species (when the number of subgroups is less than or equal to 5). (C) Box plot of Shannon’s index for *C. japonica* injected with 10^5^ CFU/L of *A. hydrophila* (AH) and control (CK).

### 3.2. Annotation and assessment of species

The distribution of the four dominant phyla was relatively similar across samples in *C. japonica* infected with *A. hydrophila*, but with different trends in abundance and variation. Figure 2 shows that there were 84 CK endemic species, 77 AH endemic species and 110 shared species. Alpha diversity indices were analysed (Figure 2B) for both richness and evenness of species composition in the 12 samples. Figure 2C shows that there were only 10 samples with a Shannon index of 2 or more for AH and CK.

### 3.3. Species-specific phylogenetic analysis

The phylogenetic evolutionary tree in Figure 3A for Polaribacter and Mesoflavibacter shows that the relative abundance of AH of Polaribacter after infection with *A. hydrophila* was 0.442326. The relative abundance of AH of Mesoflavibacter after infection with *A. hydrophila* was 0.216081. The relative abundance of Photobacterium in Firmicutes and Proteobacteria in the phylogenetic evolutionary tree was 0.421645 after infection with *A. hydrophila* and 0.288675 after infection with *Rhodobacter hydrophila*. In the phylogenetic tree, Psychroserpens in Fusobacteria had a higher AH relative abundance of 0.290992 after infection with *A. hydrophila*, and Fusibacter in Firmicutes had a higher AH relative abundance of 0.288675 after infection with *A. hydrophila*. The relative abundance of Phormidium in Cyanobacteria was 0.288675 after infection with *A. hydrophila*.

**Figure 3:**
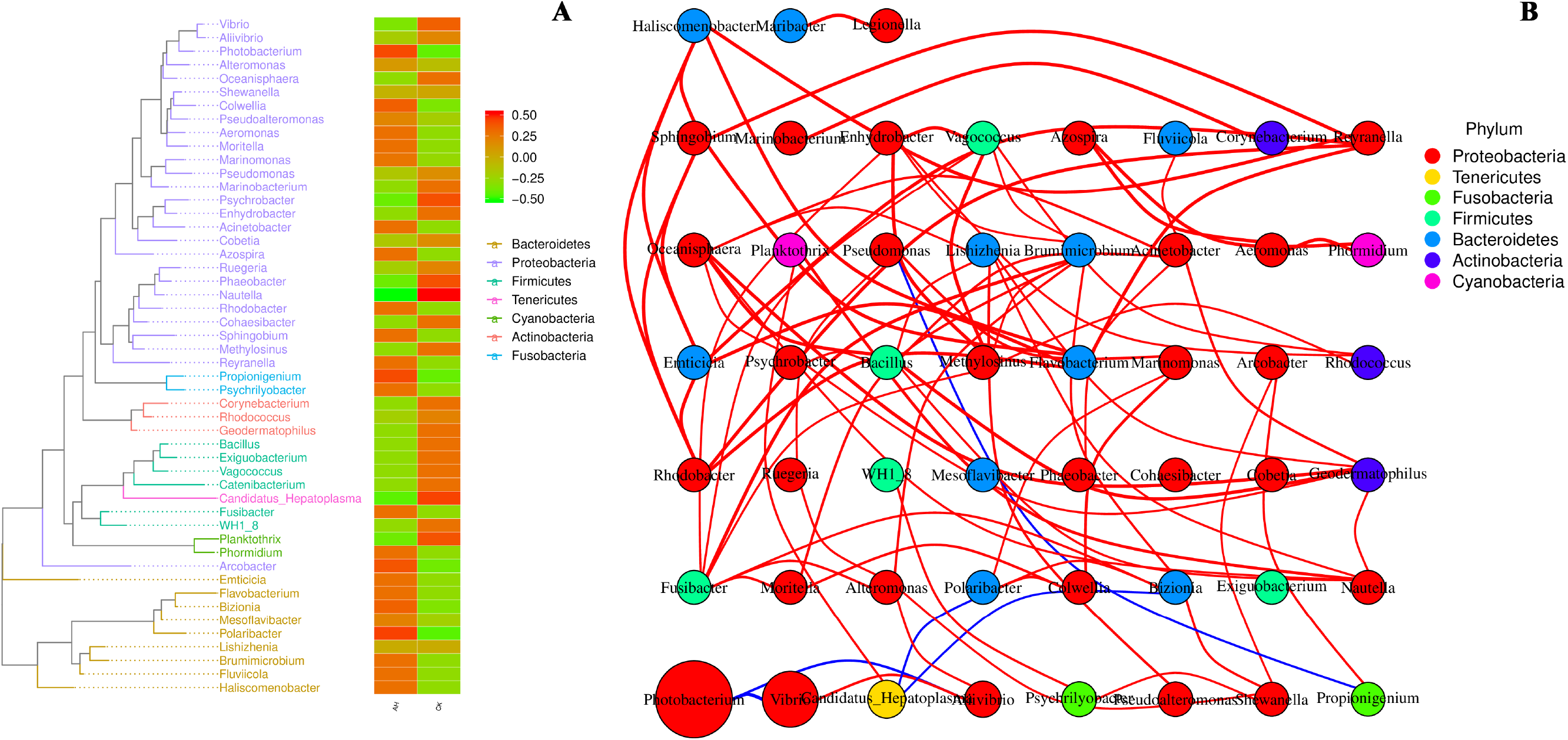
(A) Heat map of the phylogenetic evolutionary tree and the distribution of abundance between groups. The evolutionary tree is shown on the left, the different coloured branches represent different clades and each end branch represents an OTU. The end annotates the genus classification to which the corresponding OTU belongs. The heat map on the right is the normalised abundance. The mean of the abundance is 0 and the standard deviation is 0.5. Twelve samples were divided into two groups: AH and CK. (B) Species interaction network at the taxonomic level. Circles represent a species, the size represents its relative abundance, different colours represent different species phylum classifications, lines between circles represent a significant correlation between these two species (p-value less than 0.05). Red lines indicate positive correlations, blue lines indicate negative correlations. The thicker the line, the larger the absolute value of the correlation coefficient.

### 3.4. Species interaction network analysis

As can be seen in Figure 3B, Rhodobacter has a mutually supportive relationship with Emticia, Sphingobium, Bacillus, Haliscomenobacter and Reyranella. Polaribacter has a mutually reinforcing relationship with Marinomonas and Bizionia and Polaribacter, whereas Hepatoplasma has a suppressive effect. Psychrilyobacter also has a mutually reinforcing relationship with Alteromonas, Shewanella, WH1_8. Mesoflavibacter has a mutually promoting relationship with Phaeobacter, Colwellia, Bacillus, Psychrobacter and Cohaesibacter.

### 3.5. Statistical analysis of metabolites

Percentage content of 20 metabolites in the gut of *C. japonica*: Glycine in AH (0.069%), CK (0.075); Galactose in AH (0.073%), CK (0.63%); Proline in AH (0.068%), CK (0.063%); Tyrosine in AH (0.063%), CK (0.059%). Leucine in AH (0.059%), CK (0.059%); Phosphoric acid in AH (0.053%), CK (0.058%); Glucose in AH (0.061%), CK (0.050%) (Figure 4A). These findings demonstrate that there is still a direct difference between AH and CK. PLS-DA looks for factors that can find the maximum distinction between sample groupings (a factor can be interpreted as a weighted sum of all metabolites). Discriminant analysis encodes the discontinuous categorical variable to be predicted as a latent variable, which is continuous, so that a regression can be created between the explanatory and latent variables and solved using the theory of least squares regression. PLS-DA finds a linear regression model by projecting the predictor and observed variables separately into a new space (the dimensions of the new space are independent of each other and there is no covariance problem). In Figure 4B, each point corresponds to a sample and the PLS-DA effect plot is the value of the two factors that discriminate the best. The AH samples were found to be concentrated on the left side of the PLS-DA effect plot and CK on the right side of the PLS-DA effect plot.

**Figure 4:**
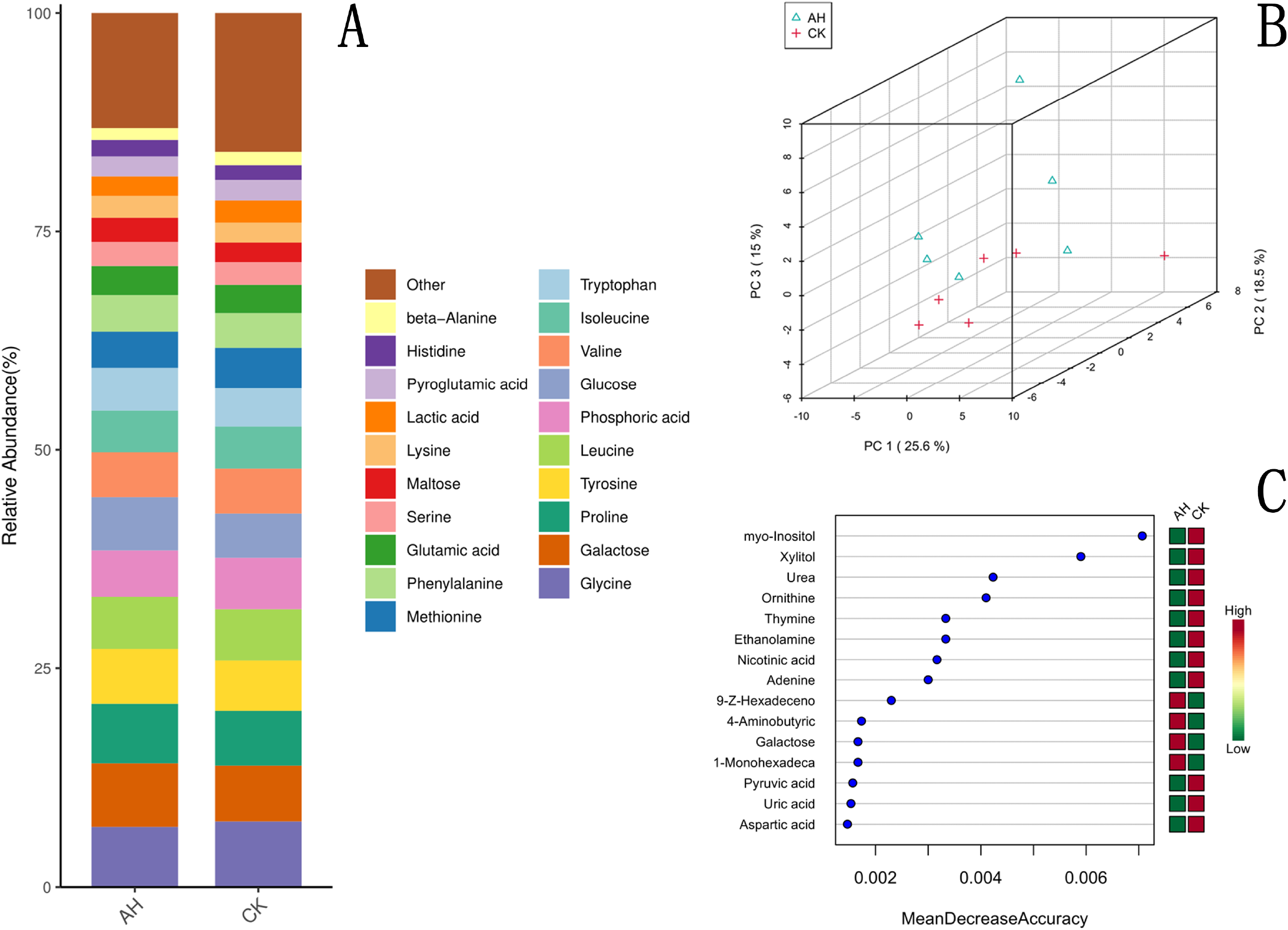
(A) Histogram of the top 20 metabolites. *C. japonica* injected with 10^5^ CFU/L of *A. hydrophila* (AH) and control (CK). (B) PLS-DA point cloud, *C. japonica* injected with 10^5^ CFU/L of *A. hydrophila* (AH) and control (CK). (C) The top 15 metabolites in the random forest. The horizontal coordinate on the left panel is “Mean Decrease Accuracy”; the right panel is a heat map of the levels of the 15 metabolites in the two subgroups. *C. japonica* injected with 10^5^ CFU/L of *A. hydrophila* (AH) and the control group (CK).

### 3.6. Metabolic pathway analysis

The degree of reduction in the predictive accuracy of the random forest by making the value of a metabolite a random number is the “Mean Decrease Accuracy”. Among the 15 most important metabolites, glucose, 9-Z-octadecenoic acid, 4-hydroxyproline and 1-monohexadecanoylglycerol increased the most in *C. japonica* infected with *A. hydrophila* (Figure 4C). The levels of xylitol, ornithine and uracil were decreased in *C. japonica* infected with *A. hydrophila*. The metabolic pathways that were significantly enriched in differential metabolites are shown in Figure 5A. Metabolites that increased in the gut of *C. japonica* after infection with *A. hydrophila* were associated with beta-alanine metabolism, nitrogen metabolism, inositol phosphate metabolism, galactose metabolism, histidine metabolism, ascorbate and aldarate metabolism, fatty acid biosynthesis, aminoacyl-tRNA biosynthesis and arginine and proline metabolism.

**Figure 5:**
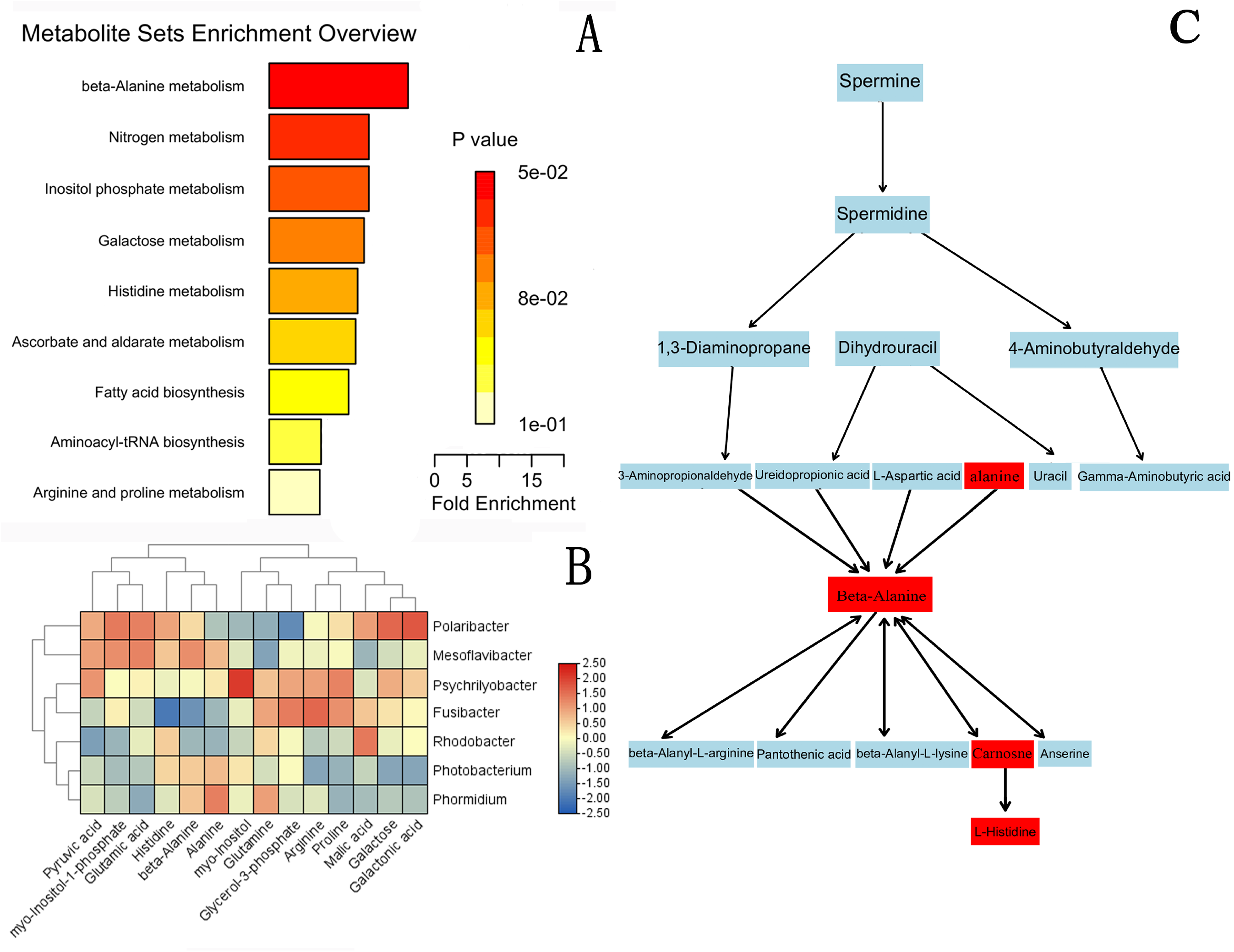
(A) RA enrichment analysis, the horizontal coordinate is the enrichment multiplier, which is the number of observed metabolites/theoretical metabolites in the metabolic pathway. The p-value magnitude is indicated by colour, the darker the colour, the smaller the p-value. (B) Association analysis results in the horizontal axis are metabolites, vertical axis are genus. Colours represent the r value, red indicates positive correlation, blue indicates negative correlation. The darker the colour the stronger the correlation; asterisks indicate that the p-value of the correlation is less than or equal to 2.50. (C) Metabolites and metabolic pathways in *C. japonica* (Acipenser japonicus) after infection with *A. hydrophila*. The red metabolites are those that differed significantly between subgroups

As can be seen in Figure 5C, alanine, beta-alanine, carnosine and L-histidine levels were increased.

### 3.7. 16S rRNA and metabolite association analysis

Polaribacter showed a positive correlation with pyruvic acid, myo-inosit-1-phosphate, beta-alanine, glutamic acid, histidine, galactose and galactonic acid. Mesoflavibacter showed a positive correlation with pyruvic acid, myo-inosit-1-phosphate, beta-alanine, glutamic acid, histidine and alanine. Psychrilyobacter showed positive correlations with alanine, myo-inositol, glycerol-3-phosphate, arginine and malic acid. Fusibacter showed a positive correlation with alanine, myo-inositol, glycerol-3-phosphate, arginine and malic acid. Rhodobacter showed positive correlations with malic acid, Glutamine and histidine (Figure 5B). Photobacterium showed positive correlations with beta-alanine, alanine and histidine. Phormidium also showed positive correlations with beta-alanine, alanine and glutamine. The activity of metabolite related enzymes such as lipid peroxides, SOD, i-NOS, GST, GSH, LPO, GPX and POX showed a downward trend as shown in Figure 8. The activity of THL, HK and PK enzymes was upregulated. The level of NO also showed a downward trend (Figure 6).

**Figure 6:**
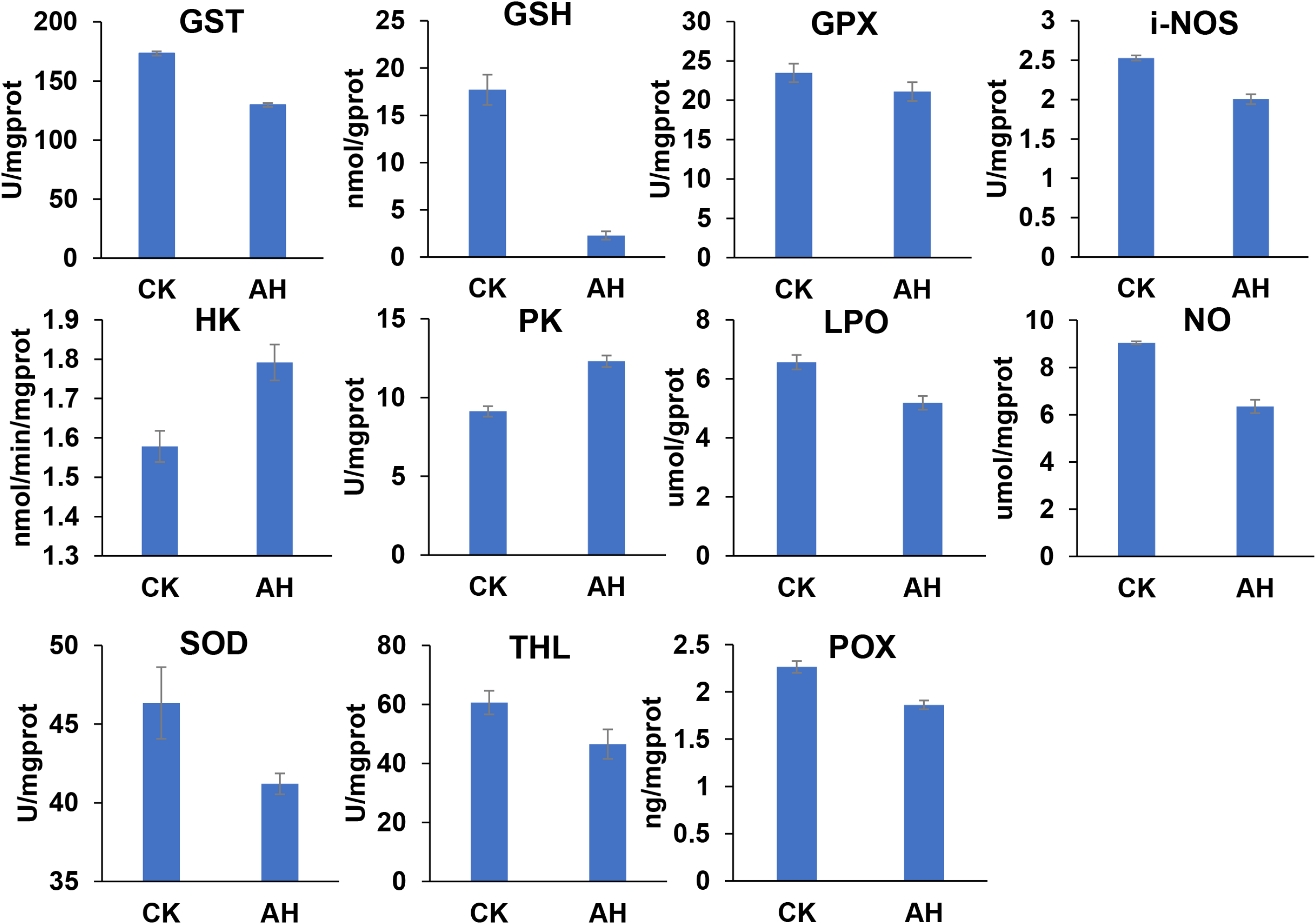
Relevant enzymes in hepatopancreas tissue of *C. japonica* treated with *A. hydrophila*. AH, the experimental group; CK, the control group. *Significantly (P < 0.05) different from control values.

## 4. Discussion

The relative abundance of some beneficial intestinal genera including Photobacterium, Rhodobacter, Polaribacter, Mesoflavibacter, Fusibacter and Phormidium was increased in *C. japonica* after infection with *A. hydrophila*. *Litopenaeus vannamei* had a significantly higher abundance of gut Photobacterium after infection with white spot syndrome virus (WSSV), which inhibits the composition and function of the gut microbiota in vivo and prevents WSSV infection (13). Research over the years has found that the most predominant species in healthy fish is Rhodobacter, and that co-culture with Roseobacter isolates and *Vibrio anguillarum* in artificial seawater-based phytoplankton media reduces the viability of *Vibrio anguillarum* by approximately 10-fold or more (14). Cultivation of Phaeobacter and Vibrio together under different growth conditions antagonised Vibrio *anguillarum*, reducing the number (15). Alkaloids also increased probiotic Rhodobacter during immunostimulatory and disease resistance in fish (16). Some bacteria in Polaribacter can degrade algal polysaccharides (17). Phormidium produces lipids such as monogalactose diacylglycerol and digalactose diacylglycerol, which are anti-inflammatory and antibacterial active substances (18). The enrichment of Mesoflavibacter in the gut of American white shrimp fed resistant starch may be involved in the degradation of toxins and the production of beneficial metabolites, and a significant elevation of Mesoflavibacter could prevent the reduction of potential pathogens (Formosa and Pseudoalteromonas) (19). The key bacterial members associated with overweight shrimps were mainly from Firmicutes (short bacilli and Fusibacter), both of which were found to be in higher abundance than those with less weight. Fusibacter in Firmicutes has been identified as a potential probiotic and antimicrobial peptide producer (20) in sea bass. From the above findings, it is evident that these increased probiotic abundances can suppress *A. hydrophila* through numerical dominance, with mechanisms that include an anti-pathogenic chemical similar to an interferon protein molecule that can be induced by bacterial molecules to be released by immune cells(21). Presumably, this mechanism can interfere with the population sensing of *A. hydrophila* in *C. japonica* pathogens through signal hijacking.

Increased metabolites in the gut of *C. japonica* after infection with *A. hydrophila* include beta-alanine metabolism, nitrogen metabolism, inositol phosphate metabolism, galactose metabolism, histidine metabolism, ascorbate and aldarate metabolism, fatty acid biosynthesis, aminoacyl-tRNA biosynthesis, arginine and proline metabolism (Figure 9). In this experiment, the metabolites of nitrogen and histidine were more abundant in the gut of *C. japonica* after being infected with *A. hydrophila*. Nitrogen is a key element in biological systems; it is a fundamental component of amino acids, proteins and nucleic acids. Most amino acids are broken down in the liver, while some can be broken down in the intestine(22). In hepatocytes, several amino acids can be degraded by specific histidases and NH_3_ is released in the cytosol (23). Aquatic animals excrete 50% of their nitrogenous waste as ammonia (24). Certain amino acids can be converted to glutamate; they include arginine, glutamine, histidine and proline. Glutamate is transaminated by alanine transaminase, which catalyses the reaction of glutamate with pyruvate to form alpha-ketoglutarate and alanine without releasing ammonia (25). The alpha-ketoglutarate produced can be directed into the tricarboxylic acid cycle and partially catabolised to malic acid. Malic acid can be directed out of the tricarboxylic acid cycle by malic enzymes and converted to pyruvate. This would provide a continuous supply of pyruvate to sustain the transamine reaction catalysed by alanine transaminase to form α-ketoglutarate and alanine (26). According to this theory, the metabolites of arginine, histidine and proline in the gut of *C. japonica* after infection with *A. hydrophila* are less than those of alanine. In this experiment, GPX, GST and GSH of the glutamate metabolic pathway were measured, and GST and GSH activities were found to be lower than in normal *C. japonica*, while GPX was not significantly changed. The activity of these enzymes showed that the glutamate metabolites were relatively low, inhibiting their redox reactions, and that the excess glutamate may have been involved in the pyruvate reaction to produce α-ketoglutarate and alanine. Metabolites in the gut of *C. japonica* infected with *A. hydrophila* are most abundant in alanine metabolites, which have two isomers, α-alanine and β-alanine. Alanine metabolites are the most important metabolites in the gut of *C. japonica* infected with *A. hydrophila*.

Myostatin is a dipeptide that is found in high concentrations in skeletal muscle. It is synthesised by the amino acids L-histidine and β-alanine, catalysed by myostatin synthase. In this experiment, we found that THL, HK and PK enzyme activities were upregulated in *C. japonica* after infection with *A. hydrophila*. Myostatin readily glycosylates with aldose and ketose, inhibits dihydroxyacetone glycosylation of histidine and Lys residues, and resists glycosylation and cross-linking of ribose, deoxyribose and fructose(27). The results of our study are quite close to those of Hipkiss et al. In this experiment, we found that lipid peroxidation was reduced in *C. japonica* after infection with *A. hydrophila* and that increased β-alanine metabolites could potentially reduce the production of free radicals in skeletal muscle and reduce lipid peroxidation. It was found that the hepatopancreas NO content was reduced and nitric oxide synthase (i-NOS) activity was also reduced after infection with *A. hydrophila* and that this increased NH_3_ content in nitrogen metabolism and water reduced blood cell counts and haemocyanin levels, inhibited blood cell phagocytosis, reduced SOD and phenoloxidase activity, and disrupted immune defence systems, including immune cells and immune response factors(28). Increased NH_3_-N in water is reported to reduce the immunity of crabs to pathogens and their ability to scavenge free radicals(29). In the present experiment, *C. japonica* showed reduced activity of SOD and phenoloxidase in the hepatopancreas after infection with *A. hydrophila*.

The literature that has investigated the interaction between probiotics and pathogenic bacteria support the idea that probiotics inhibit pathogenic bacteria by competing for adhesion sites (30), aggregating with pathogenic bacteria and producing metabolites (31). However, the presence of conditionally pathogenic bacteria in the gut does not disappear with the presence of intestinal probiotics, suggesting that there may be interactions between probiotics and pathogenic bacteria other than antagonism(32). Therefore, we found that Mesoflavibacter, a representative of the intestinal probiotics of *C. japonica*, had a mutually promoting relationship with Phaeobacter, Colwellia, Bacillus, Psychrobacter and Cohaesibacter; Rhodobacter with Emticia, Sphingobium, Bacillus, Haliscomenobacter and Reyranella; and Polaribacter with Marinomonas and Bizionia (Figure 5). Kernbauer et al. (32) reported that some pathogens cause diarrhoea and can promote enterocyte colonization, which helps to restore structural and functional damage to the intestine caused by enteritis. As a typical conditional pathogen, *L. monocytogenes* is present in the intestine of many healthy individuals (33). To date, the biological significance of its presence has not been reported. This suggests that conditional pathogenic bacteria in the gut of healthy crabs may also have a symbiotic relationship with intestinal probiotics, promoting their growth and multiplication and enhancing their prebiotic effect. This study verifies that Phormidium interacts with Aeromonas and Azopira, and it can be inferred that *Aeromonas hydrophila*, a pathogenic bacterium in *Aeromonas*, can also promote the growth of other probiotics. It is hoped that more pathogens and probiotics will be selected for further study to examine this stimulatory effect and molecular mechanism and to explain this phenomenon more clearly.

Different tissues of animals, such as the skin, the hepatopancreas and the intestine, contain a large number of bacteria. Due to improvements in 16s sequencing technology, tens of millions of different microbial genes have been identified in the human gut, thus exceeding the number of other animal genomes by a factor of several hundred (34). This huge number of genes signifies that bacteria have a greater metabolic function in their hosts, synthesising essential amino acids and potentially participating in the metabolism of the host organism(35). As a result, the *C. japonica* gut microbiome is considered to be a tissue with the metabolic potential to influence the metabolism of *C. japonica*. This is because the gut microbiome allows the nutrients contained in food to be processed and allows their components to be adsorbed and reused by crab cells. Thanks to improved methods of metabolite (metabolomics) analysis, recent analyses have shown that many of these metabolites originate from the metabolism of intestinal bacteria. Classes such as Mesoflavibacter showed positive correlations with pyruvic acid, myo-inosit-1-phosphate, beta-alanine, glutamic acid, histidine and alanine; Fusibacter showed positive correlations with alanine, myo-inositol, glycerol-3-phosphate, arginine and malic acid; Rhodobacter showed a positive correlation with malic acid, glutamine and histidine; Photobacterium showed a positive correlation with beta-alanine, alanine and histidine; Phormidium showed a positive correlation with beta-alanine, alanine and glutamine. This suggests that metabolites increased with the abundance of bacteria and therefore showed a positive correlation. It is possible that the autoimmune system and the entry of *A. hydrophila* into the intestinal tissues of *C. japonica* react immunologically and that the organism is producing certain metabolites which may be metabolised by various bacteria and absorbed into the circulation. In addition, some of these metabolites are modified or bound in the liver to produce microbiota-host co-metabolites(35). These microbial metabolites or co-metabolites can resist invasion by *A. hydrophila*. For example, through specific receptors such as short-chain fatty acids, indoles or myostatin, as described previously(36).

## 5. Conclusion

In conclusion, the results of this study suggest that some beneficial genera of bacteria in the intestine of *C. japonica* can inhibit *A. hydrophila* after infection. The most abundant metabolites in the gut of *C. japonica* infected with *A. hydrophila* were alanine metabolites. In this experiment, lipid, peroxide, SOD, i-NOS, phenol oxidase, GST and GSH and NO levels were found to be decreased in *C. japonica* infected with *A. hydrophila* in the intestine. The enzymatic activity of THL, HK and PK was upregulated in *A. hydrophila*-infected *C. japonica*. The probiotics in the intestinal tract of *C. japonica* have a mutually reinforcing relationship, promoting their growth and reproduction and enhancing their prebiotic effect. *C. japonica* metabolites showed a positive correlation with probiotic flora, suggesting that metabolites increased with the abundance of bacteria and that microbial metabolites or co-metabolites could resist invasion by *A. hydrophila*. The above results may contribute to further studies investigating the resistance of the crab to invading pathogenic bacteria. They also provide information for future research on relevant probiotic and metabolite pathways.

**Table 1.** Statistical table of DADA2 denoising to generate (Operational Taxonomic Units) OTUs.

| sample-id | input | filtered | denoised | merged | non-chimeric |
| --- | --- | --- | --- | --- | --- |
| AH1 | 51820.0 | 46995.0 | 46995.0 | 46262.0 | 42540.0 |
| AH2 | 53801.0 | 48149.0 | 48149.0 | 44588.0 | 27026.0 |
| AH3 | 40419.0 | 37184.0 | 37184.0 | 34778.0 | 16628.0 |
| AH4 | 48431.0 | 43146.0 | 43146.0 | 41123.0 | 33171.0 |
| AH5 | 58952.0 | 53367.0 | 53367.0 | 49501.0 | 25823.0 |
| AH6 | 49210.0 | 44315.0 | 44315.0 | 43513.0 | 40142.0 |
| CK1 | 52331.0 | 47532.0 | 47532.0 | 43157.0 | 21781.0 |
| CK2 | 48898.0 | 44245.0 | 44245.0 | 40835.0 | 22046.0 |
| CK3 | 38808.0 | 34699.0 | 34699.0 | 33159.0 | 26160.0 |
| CK4 | 43823.0 | 39674.0 | 39674.0 | 38840.0 | 27073.0 |
| CK5 | 58821.0 | 53665.0 | 53665.0 | 50405.0 | 27483.0 |
| CK6 | 32453.0 | 28733.0 | 28733.0 | 26260.0 | 14226.0 |

## 6. Acknowledgements

The authors would like to thank for Mengling Sun her help of sampling. This work was supported by National Key R&D Program of China (2020YFD0900305) Science Foundation of Jiangsu (NO. BK20191488) in China, Major project of hydrobios resources in Jiangsu province (ZYHB16-3) and agricultural major new variety creation project in Jiangsu province (PZCZ201743).

